# Consistent pre-stimulus influences on auditory perception across the lifespan

**DOI:** 10.1101/378851

**Authors:** Steven W. McNair, Stephanie J. Kayser, Christoph Kayser

## Abstract

As we get older, perception in cluttered environments becomes increasingly difficult as a result of changes in peripheral and central neural processes. Given the aging society it is important to understand the neural mechanisms constraining perception in the elderly. In young participants, the state of rhythmic brain activity prior to a stimulus has been shown to modulate the neural encoding and perceptual impact of this stimulus – yet it remains unclear whether, and if so, how, the perceptual relevance of pre-stimulus activity changes with age. Using the auditory system as a model, we recorded EEG activity during a frequency discrimination task from younger and older human listeners. By combining single-trial EEG decoding with linear modelling we demonstrate consistent statistical relations between pre-stimulus power and the encoding of sensory evidence in short-latency EEG components, and more variable relations between prestimulus phase and subjects’ decisions in fronto-parietal EEG components. At the same time, we observed a significant slowing of auditory evoked responses and a flattening of the overall EEG frequency spectrum in the older listeners. Our results point to mechanistically consistent relations between rhythmic brain activity and sensory encoding that emerge in large despite changes in neural response latencies and the relative amplitude of rhythmic brain activity with age.

## 1. Introduction

In everyday life our acoustic environments are often teeming with incoming information. Yet, the auditory brain manages to filter target information from noise seamlessly, at least in the young and healthy brain (Bregman, 1994). With advancing age listening becomes more challenging, particularly in “cocktail party” scenarios (de Villers-Sidani et al., 2010; Pichora-Fuller et al., 2017; Rossi-Katz and Arehart, 2009). This difficulty could arise from age-related changes in peripheral and central auditory processes (Anderson et al., 2013; Clinard et al., 2010; Clinard and Cotter, 2015; Harris and Dubno, 2017), such as the poorer encoding in early sensory regions (Grose and Mamo, 2012; He et al., 2007; Mahajan et al., 2017; Paraouty et al., 2016; Wallaert et al., 2016). Changes in higher cognitive processes may also influence older adults’ performance via top-down feedback (Henry et al., 2017), through reduced attentional flexibility (Nunez et al., 2015; Zanto and Gazzaley, 2014), or changes in decision criteria when reporting perceptual performance (Dully et al., 2018).

As shown by recent work perception depends not only on the qualities of the sensory signal but also on the state of the brain prior to stimulus occurrence (Henry et al., 2017, 2014; Henry and Obleser, 2012; Kayser et al., 2016; Ng et al., 2012; Pinheiro et al., 2017). In many studies, the state (power or phase) of pre-stimulus rhythmic brain activity has been predictive of perceptual performance in a variety of tasks, in line with the view that perception in general is controlled by a cascade of rhythmic neural processes (Schroeder et al., 2010; VanRullen, 2016). Furthermore, changes in top-down influences by attentional and cognitive strategies are also reflected in rhythmic brain activity, especially in the alpha and beta bands (Henry et al., 2017; Petersen et al., 2015; Strauss et al., 2015; Wöstmann et al., 2017). In this context of relating rhythmic brain activity to perception we recently described two putative mechanisms by which pre-stimulus activity shapes auditory perceptual decisions in younger adults (Kayser et al., 2016): in that study the power of low-frequency and beta activity affected the encoding of sensory information in early auditory regions, while the phase of the alpha band influenced decision processes in high-level regions.

This importance of rhythmic activity for perception raises the question as to whether the underlying mechanisms and relevant time scales are conserved across the age span. For example, it is known that cognitive and neural processes become slower with age (Bieniek et al., 2013; Price et al., 2017; Salthouse, 1996), which is reflected in changes in the amplitude and latency of auditory evoked responses (Harris et al., 2008; Henry et al., 2017; Tremblay et al., 2003), an increase in response stereotypy (Garrett et al., 2013, 2011; Herrmann et al., 2016), and changes in the slope of the overall frequency spectrum of brain activity (Hong and Rebec, 2012; Tran et al., 2016; Voytek et al., 2015). This makes it possible that the patterns of rhythmic brain activity that shape perception systematically change with age.

We here capitalized on our previous study in a group of younger subjects to directly probe whether the mechanisms linking pre-stimulus brain activity, sensory encoding and decision-making are conserved with age. Specifically, we compared behavioural and electroencephalogram (EEG) data from younger (<30 years) and older (>65 years) listeners obtained during an auditory frequency discrimination-in-noise task. For each group we linked pre-stimulus oscillatory activity to neural signatures of stimulus encoding and decision making using single trial modelling. We expected to observe the same patterns of statistical relations between neural activity, sensory encoding and choice in both groups (i.e. significant relations between the same variables), but with the possibility that the precise time scales (i.e. frequency bands of brain activity) differed. For comparison, we also quantified age-related changes in the amplitude and timing of evoked responses and the spectral slope of the overall EEG signal.

## 2. Materials and Methods

### 2.1. Participants

We collected data from 16 younger (6 male; mean ± SD age, 23.9 ± 1.1 years) and 17 older adults (8 male; mean ± SD age, 68.4 ± 3.6 years). We have reported data from the younger group, with the exclusion of PSD and AEP analyses, in our previous study Kayser et al. (2016) (the frequency task there). For this reason, we had set the target sample size for the group of older subjects to match the size of the younger group. Younger participants had normal self-reported hearing, as measured by the Better Hearing Institute Quick Hearing Questionnaire (Kochkin and Bentler, 2010). Older participants had no more than mild hearing loss as measured by the Better Hearing Institute Quick Hearing Questionnaire, Tinnitus Handicap Inventory (THI, where applicable; McCombe et al., 2001) and pure-tone audiometric (PTA) procedures. The PTA procedure was presented via MATLAB (2015b; The MathWorks Inc., Natick, MA) and was designed in accordance with guidelines from the British Society of Audiology (BSA, British Society of Audiology, 2012). We tested participants’ hearing thresholds at frequencies of 250Hz, 500Hz, 1000Hz, 2000Hz, 4000Hz and 8000Hz individually for each ear. Sound levels were calibrated using a Bruel&Kjaer sound-level meter. Older participants were also screened for cognitive impairment using the Montreal Cognitive Assessment (MoCA, Nasreddine et al., 2005), D2 test of attention (Brickenkamp and Zillmer, 1998), and the digit span working memory test (Turner & Ridsdale, 2004). Due to possible variability in participants’ frequency discrimination abilities (Foxton et al., 2009; Liang et al., 2016; Semal and Demany, 2006), frequency difference limens (see below) were tested both at screening and immediately prior to the main experiment for each group. Group-level auditory and cognitive test scores are shown in Table 1. Four older participants were excluded at screening based on pre-defined criteria: two participants had moderate to severe hearing loss, as indicated by PTA testing, and in two participants frequency difference limens could not be measured reliably.

**Table 1.**
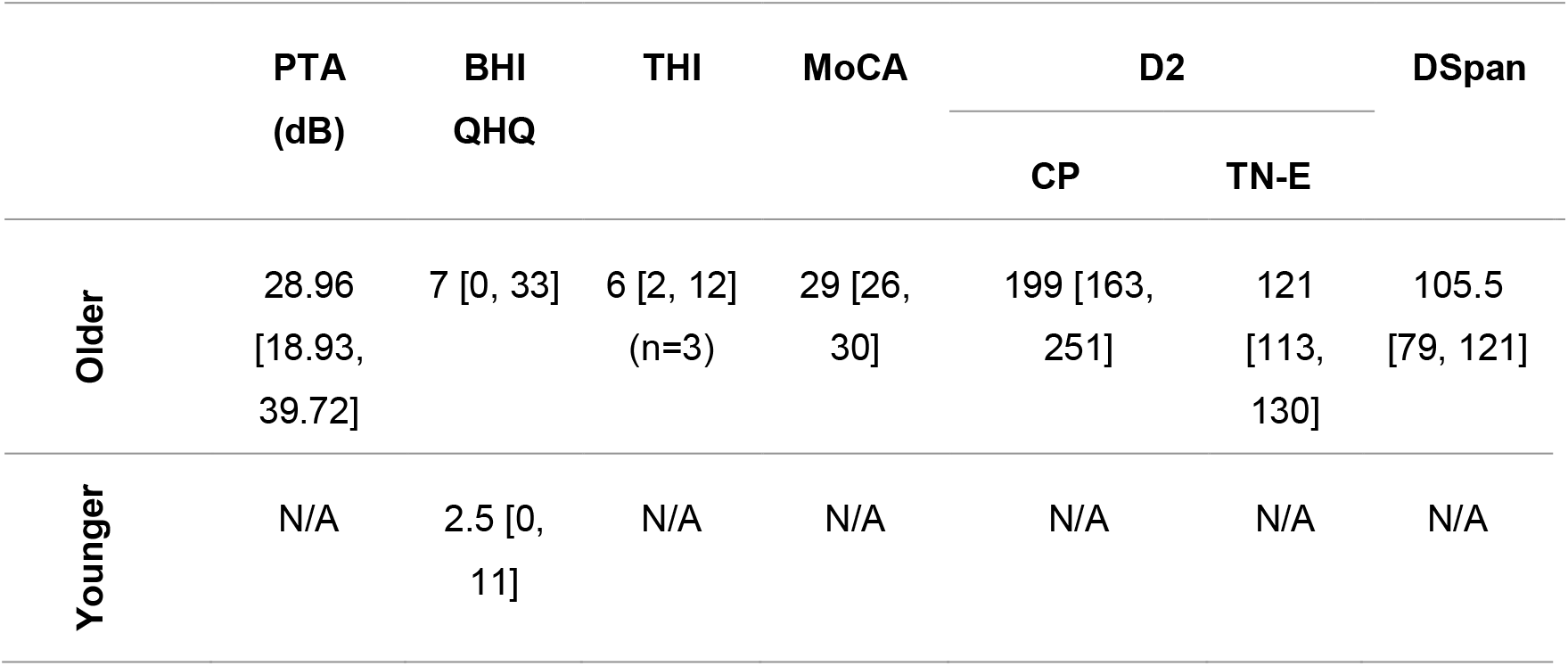
Auditory and cognitive test scores. Screening scores for younger (where applicable) and older participants who passed screening. Hearing scores are derived from pure tone audiometry (PTA), Better Hearing Institute Quick Hearing Questionnaire (BHI QHQ), and Tinnitus Handicap Inventory (THI). THI was administered only as applicable, thus n is reported. Cognitive test scores are derived from Montreal Cognitive Assessment (MoCA), D2 test of Attention (D2) and digit span (DSpan) tests. Scores correspond to median across all participants in each age group. Square brackets indicate minimum and maximum scores. N/A indicates where data was not available.

Participants indicated no history of mental/neuropsychological disorders, stroke, or brain or ear injuries. Participants gave written informed consent and received £6/hour payment plus travel expenses for participating. This study is in accordance with the Declaration of Helsinki and was approved by the local ethics committee (College of Science and Engineering, University of Glasgow).

### 2.2. Auditory Stimuli

Participants completed a 2-alternative forced-choice auditory frequency discrimination task, as described in Kayser et al. (2016). Participants were presented with two sequential target tones embedded within a noisy background and had to discriminate which tone was higher in frequency (see Fig. 1A). Targets were pure-tones of 50ms duration (including a 5ms cosine on/off ramp) and spaced 50ms apart. Both tones were equated in intensity at a signal-to-noise-ratio of ±2dB relative to background intensity, based on the r.m.s. level. The second tone was kept at a constant 1024Hz while the first varied pseudo-randomly over 7 (younger participants) or 5 (older participants) equally-spaced (on an octave scale) levels of a frequency difference above or below the second (pseudorandomized and balanced across all trials), ranging from 0Hz difference to 2Δ in younger and 2.5ΔHz in older participants (where Δ is the participants’ own 70% correct frequency difference limen). The reason we reduced the number of frequency levels for the older adults was to keep the experimental duration to a minimum to avoid fatigue.

**Fig. 1:**
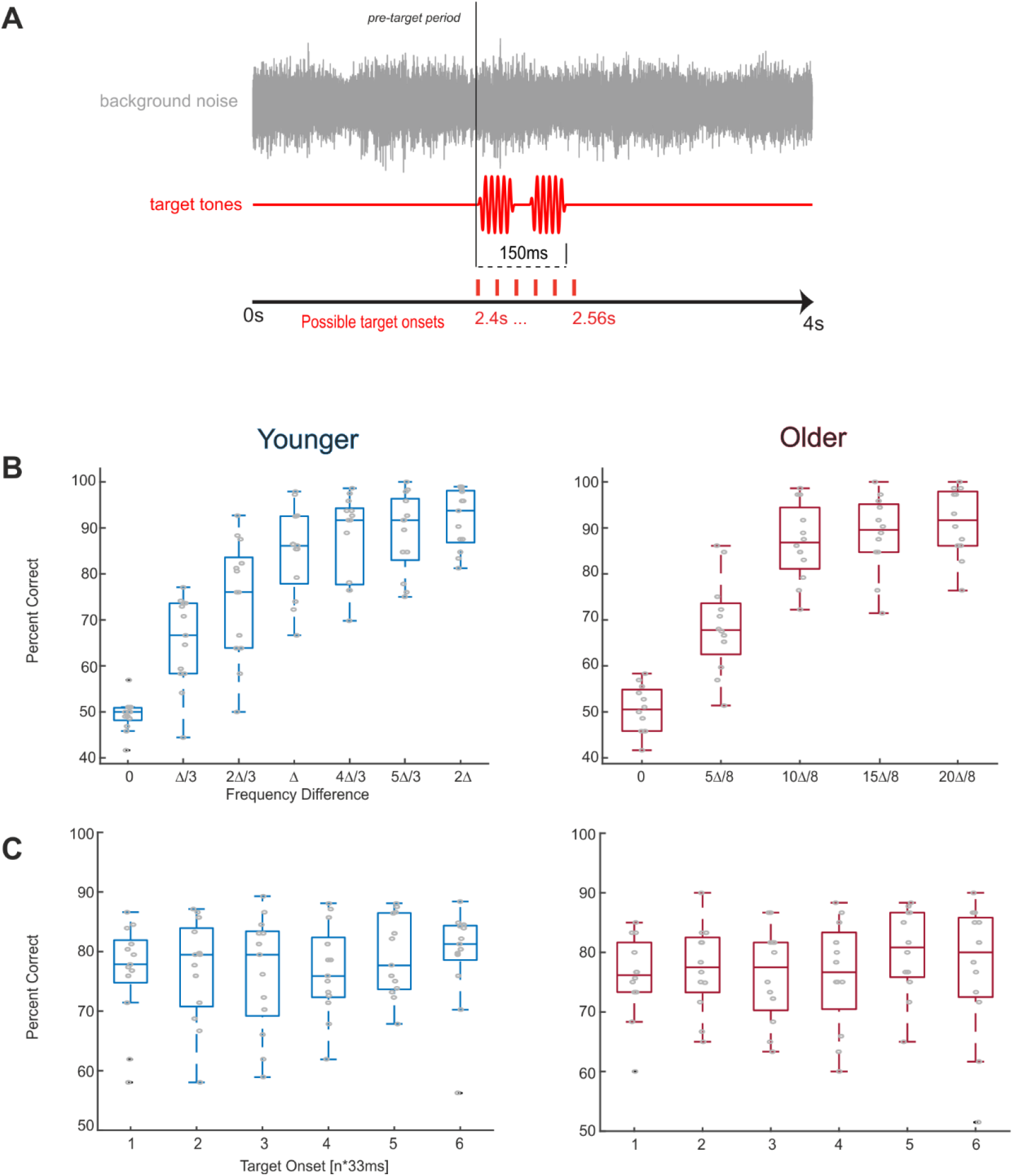
Auditory paradigm and task performance. (A) Auditory paradigm. Pure tone targets (50ms duration, spaced 50ms apart), were presented at one of six possible onsets against a continuous background noise cacophony. The second tone was kept at 1024Hz while the first varied over 7 (younger adults) or 5 (older adults) levels of frequency difference, titrated around participants’ own frequency difference limens, Δ. (B) Group level task performance as a function of frequency difference level, averaged across target positions. Younger and older adults show comparable task performance. (C) Group level task performance as a function of target position, averaged across frequency difference levels. There were no significant effects of target position on performance in either group and overall there was no significant difference between groups (across frequency levels and target positions). Grey circles indicate individual subject data.

### 2.3. Experimental Procedure

Auditory stimuli were controlled using MATLAB using the Psychophysics Toolbox Version 3 (Brainard, 1997) and presented using Sennheiser headphones. Prior to the main experiment, participants completed training trials to familiarize themselves with the task and their frequency difference (in noise) limens were obtained using three interleaved 2-down-1-up staircase procedures. In the actual experiment target tones were presented at one of six possible pseudorandom delays (2400 + n*33ms, where n = 0 … 5) relative to background onset. Trials were separated by an inter-trial period uniformly distributed between 1700-2200ms, and were presented in a block design. Each block contained 120 trials with each participant completing 360 trials in total.

### 2.4. EEG Recording and Pre-processing

EEG signals were recorded in a dark and electrically-attenuated room using an active 64-channel BioSemi system (BioSemi B.V., Netherlands). Electrooculogram (EOG) was derived from four electrodes placed at the outer canthi and below each eye. Electrode offsets were kept below 25mV and data were recorded at a 500Hz sampling rate using a 208Hz low-pass filter.

Pre-processing and data cleaning was carried out as described previously in Kayser et al. (2016). In brief, the data were filtered between 1-70Hz and Independent Components Analysis was used to remove eye movement and blink artefacts (Debener et al., 2010) and muscle artefacts (Beirne and Patuzzi, 1999; Hipp and Siegel, 2013). Trials were rejected if the peak signal on any electrode exceeded ± 100 μV or if participants responded faster than 400ms following the first target tone. Based on these criteria we rejected an average of 5% of trials. EEG signals were re-referenced to the common average for further analysis.

### 2.5. Analysis methods

#### 2.5.1. Evoked Responses

We compute ERPs in response to the onset of the noisy acoustic background based on trial-averaged data over a 3×3 grid of central channels (FC1, FCz, FC2, C1, Cz, C2, CP1, CPz, CP2). Individual participants’ P1, N1 and P2 component median peak latencies and amplitudes were then extracted and compared between groups.

#### 2.5.2. Pre-stimulus Power Spectra

We computed power PSD estimates using Welch’s method averaged across all channels in a pre-target time window of −0.6s to 0s relative to target tone onset. PSD estimates were normalized by removing individual participants’ mean PSD from their own spectra and were fit using a linear regression models to extract PSD slopes over frequencies between 1Hz and 25Hz whilst excluding alpha power between 7 and 14Hz ((Tran et al., 2016; Voytek et al., 2015).

#### 2.5.3. Single Trial Decoding of EEG Signals

To link pre-stimulus activity with perception we used the same statistical modelling approach as in our previous study (Kayser et al., 2016). We computed pre-stimulus activity in task-relevant EEG components extracted using multivariate linear discriminant analysis (Boyle et al., 2017; Kayser et al., 2016; Parra et al., 2005; Philiastides, 2006; Ratcliff et al., 2009). We searched for discriminant components within the EEG data that best discriminated between the frequency conditions (i.e. 1^st^ or 2^nd^ tone higher in frequency; see Kayser et al., 2016). This yields a one-dimensional projection, *Y*(*t*), of the EEG data, *x*(*t*), as defined by spatial weights, *w*(*t*), and a constant, *c*, as follows:

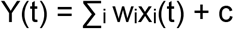

with *i* summing across channels. Classification was based on regularized linear discriminant analysis (Philiastides et al., 2014), which was applied to the EEG data in 80ms sliding windows. We assessed classifier performance using the receiver operator characteristic (ROC; referred to herein as Az) based on 10-fold cross validation. The statistical significance of the performance was assessed by shuffling condition labels 1000 times, computing the group-average Az value for each randomization, and taking the maximal Az value along time to correct for multiple comparisons (Nichols and Holmes, 2003). We estimated the corresponding forward model for each component by computing the normalized correlation between the discriminating projection and the original EEG data (Parra et al., 2005).

To select scalp projections that reflect EEG activity that was temporally consistent across subjects, we selected three systematically different components which corresponded to three continuous time windows using K-means clustering based on component topographies (see Kayser et al., 2016 for details). For each participant, we then extracted the weight (w) from the time point associated with the maximal Az value within each component for further analysis, which allowed us to incorporate between-subject variability in response timing in the analysis.

Since *Y(t)* is indicative of the extent of separability between frequency levels, we exploit this as a measure representing the amount of encoded sensory evidence about the task relevant tones (Grootswagers et al., 2017; Guggenmos et al., 2017). We computed each components’ time course by applying the respective weight to all trials and time points, resulting in a one-dimensional projection of single-trial task-related activity which we then analysed further.

#### 2.5.4. Pre-target Time-Frequency Analysis

Time-frequency representations (TFRs) were calculated using Morlet wavelets in FieldTrip (Oostenveld et al., 2011). Frequencies ranged from 2Hz to 40Hz in linear steps of 1Hz below 16Hz and 2Hz above. To achieve greater frequency smoothing at the higher frequencies the width of individual wavelets scaled with frequency (min = 4 cycles, max = 9 cycles). TFRs were calculated between −0.6s and −0.1s relative to target onset in 50ms bins. To avoid post-target contamination, we set the post-target period to zero for TFR analysis by applying a 40ms Hanning window to the last 40ms of the pre-stimulus period (Henry et al., 2014). Power was z-scored within participants and frequency bands across time and trials.

#### 2.5.5. Statistical Analyses

Group-level psychometric curves were computed for the percentage of correct responses as a function of frequency difference (averaging over temporal positions), and as a function of temporal position (averaging over frequency difference). The median performance, averaging across frequency difference levels and temporal positions, between age groups was compared using a Wilcoxon rank sum test, with effect size (r) calculated by dividing the Z value by the square root of N, where N represents the number of observations (Field, 2013). To test whether performance differed as a function of temporal position we used a non-parametric, one-way repeated-measures analysis of variance by ranks (Friedman Test).

ERP peak amplitudes/latencies and PSD slopes were compared between age groups using a non-parametric Wilcoxon rank-sum tests, with effect sizes (r) calculated following Field (2013).

To investigate the relationship between single-trial pre-stimulus activity (power/phase in particular frequency bands and time bins), sensory evidence, *Y*(*t*), extracted from each component), and perceptual choice we used linear regression modelling (Fig. 4). Model 1 tested whether pre-stimulus power/phase influences choice using regularized logistic regression. Model 2 tested whether pre-stimulus power/phase influences sensory evidence *Y(t)* using linear regression. Finally, we tested for possible mediation effects, where pre-stimulus activity state influence choice through mediation of sensory evidence (i.e. an indirect influence of pre-stimulus state on choice; see Kayser et al., 2016) using model 3: regularized logistic regression of choice on both *Y* and power/phase. Mediation effects were tested by comparing models 2 and 3. We calculated each model separately for power and phase, and for each pre-target time-frequency point. For regressions involving sensory evidence, we coded *Y*(*t*) as an unsigned variable and Z-scored it within each stimulus level, to reflect the amount of evidence about the respective stimulus. For phase, both sine- and cosine-transformed phase angles were submitted to the regression model. Mediation effects were defined by the difference in regression parameters between models 2 and 3, adjusting for dichotomous outcomes (MacKinnon et al., 2007).

Group-level statistical testing was performed using cluster-based permutation procedures and correcting for multiple comparisons across relevant dimensions, as described previously (Kayser et al., 2016). Specifically, we used 1000 randomization realizations, a 5^th^ percentile cut-off to define significant clusters, defining clusters by at least four significant neighbours, and using the cluster mass index. A two-sided test at p<0.05 was performed on the clustered data and we corrected for multiple comparisons across regression models and components using the false discovery rate (FDR) at p<0.05. We report effect sizes for clustering statistics as the cluster mass across all bins within a cluster (*T_sum_*).

The peak effect frequencies were compared across groups using a percentile bootstrap test (using 2000 samples). We randomly assigned participants to either group and compared the actual difference in group-level peak frequencies extracted from the respective statistical contrast for each regression factor to the distribution of differences in the randomized data. For this analysis effects were averaged over time for the duration of the respective clusters. Given that there were two significant clusters linking power to sensory evidence, we constrained the frequency range for the alpha/beta cluster to 8-26Hz, and the range for the low-frequency cluster to 2-13Hz in order to be able to separate these effects.

To link changes in EVP amplitudes and latencies to the peak frequencies of prestimulus effects we first computed leave-one-out estimates of the respective peak-frequencies of the pre-stimulus effects and of EVP amplitudes and latencies. We relied on a leave-out-one (Jacknife) approach as peak frequencies for pre-stimulus effects were more robust at the group-level than for individual subjects. We then used the six EVP characteristics (c.f. Fig. 2) as predictors for the peak frequency of the pre-stimulus effect across the full sample of younger and older participants in a linear regression model, for which we obtained the overall model performance and significance

**Fig. 2.**
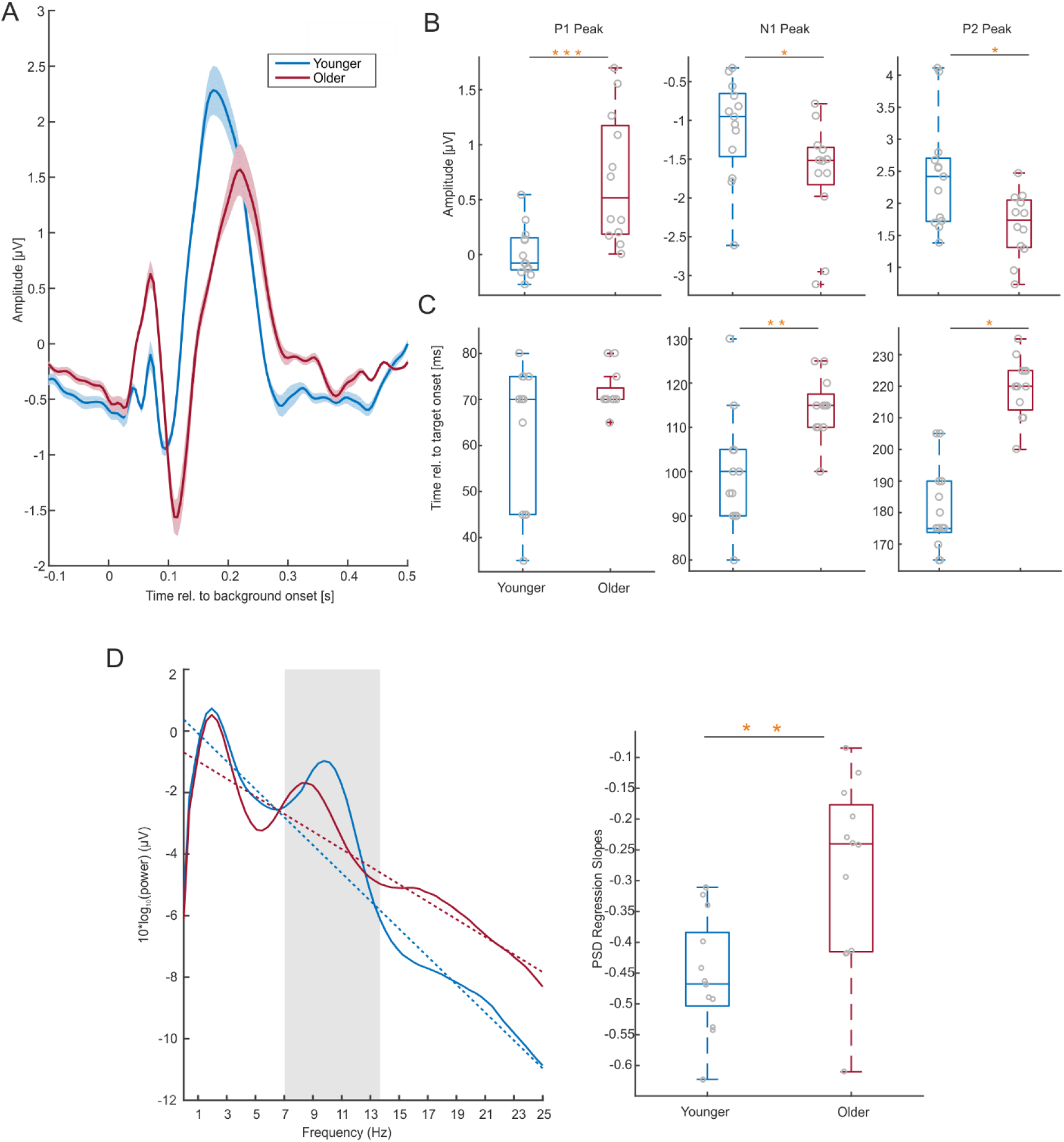
Auditory evoked responses to background onset and pre-stimulus power spectral density. (A) Grand-average AEPs over central channels (FC1, FCz, FC2, C1, Cz, C2, CP1, CPz, CP2). Both younger and older subject display a clear P1-N1-P2 potential. (B) A comparison of ERP peak amplitudes between groups revealed an age-related enhancement of P1 and N1 peaks, and a reduction in the P2 peak. (C) Component peaks were also compared in terms of latencies, revealing an age-related delay in N1 and P2 peaks. (D left panel) Group-averaged PSD estimates (smooth curves) and fitted regression slopes (dashed lines) for frequencies up to 25Hz, averaged over all channels. Slopes were computed whilst ignoring alpha power between 7-14Hz (indicated by shaded area). (D right panel) PSD slopes were reduced for the older adults. Grey circles indicate individual subject data. Yellow asterisks indicate significance as follows: * p<0.05, ** p<0.01, *** p<0.001.

## 3. Results

### 3.1. Behavioural Performance

As expected given the experimental design the overall performance was comparable across group (averaged over frequency difference level and temporal position younger median = 78.9% correct, older median = 74.4%, Z= 1.251, p = 0.211, r = 0.25; Fig. 1B). To avoid expectancy effects, target tone pairs were presented at six temporal positions relative to background onset. Friedman’s tests revealed no effect of target position on performance in either group (younger adults: X^2^(5)=5.28, p = 0.382; older adults: X^2^(5) = 8.3, p = 0.141; Fig. 1C), suggesting that any influence of pre-target activity on performance would occur without explicit cortical entrainment to the acoustic noise in either group (Henry & Obleser, 2012; Ng et al., 2012).

### 3.2. Age-related changes in auditory evoked responses

To confirm previous reports of an age-related slowing of sensory-evoked activity we compared the latency and amplitude of evoked components (P1, N1 and P2; Fig. 2A). Peak amplitudes were significantly stronger for P1 and N1 in the older group, while P2 amplitudes were reduced (P1: younger median = −0.077μV, older median = 0.517μV, Z =-3.291, p< 0.001, r = −0.658; N1: younger median = −0.95μV, older median = −1.519μV, Z=2.094, p = 0.0362, r = 0.419; P2: younger median = 2.418μV, older median = 1.736μ, Z= 2.366, p = 0.018, r = 0.473; Fig. 2B). The latencies of N1 and P2 in the older adults were significantly delayed (N1: younger median = 0.1s, older median = 0.115s, Z=-2.99, p = 0.003, r = −0.598; P2: younger median = 0.175s, older median = 0.22s, Z= −4.124, p<0.05, r = −0.825; Fig. 2C). There was no significant difference in P1 latency (younger median = 0.07s, older median = 0.07s, Z = −1.013, p = 0.311, r = −0.203).

### 3.3. Pre-stimulus PSD flattens with age

Given previous reports of changes in the power spectra of ongoing brain activity with age (Klimesch, 1999; Tran et al., 2016; Voytek et al., 2015), we analysed the spectral slope of the EEG signal (Fig. 2D). The PSD slopes of the older group were significantly flatter than those of the younger participants (younger median = −0.478dB, older median −0.24dB, Z = −2.91, p = 0.005, r = −0.582).

We also tested whether, across subjects, the observed changes in ERP latency and amplitude correlated with changes in spectral slope. Differences in PSD slope correlated significantly with differences in ERP latency for the P2 component (spearman rank-correlation: r=0.42, p=0.033, reduced slope corresponding to longer latency) but not the other ERP components (N1: r=0.016, p=0.93, P2: r=0.165, p=0.43). Differences in PSD slope also correlated with the amplitudes of the P1 (r=0.43, p=0.03) and N1 (r= −0.45, p=0.025) peaks, with a flatter PSD spectrum correlating with stronger evoked responses. There was no correlation with the P2 amplitude (r=-0.25, p=0.22).

### 3.4. Single Trial Decoding of EEG Signals

Using single-trial modelling we extracted EEG components that maximally differentiated between the stimulus conditions on which the participants task relied (1^st^ or 2^nd^ tone higher). For both groups, classification performance became significant around 0.2s following target onset (randomization test, p < 0.01, corrected for multiple comparisons along time, Fig. 3 main).

**Fig. 3.**
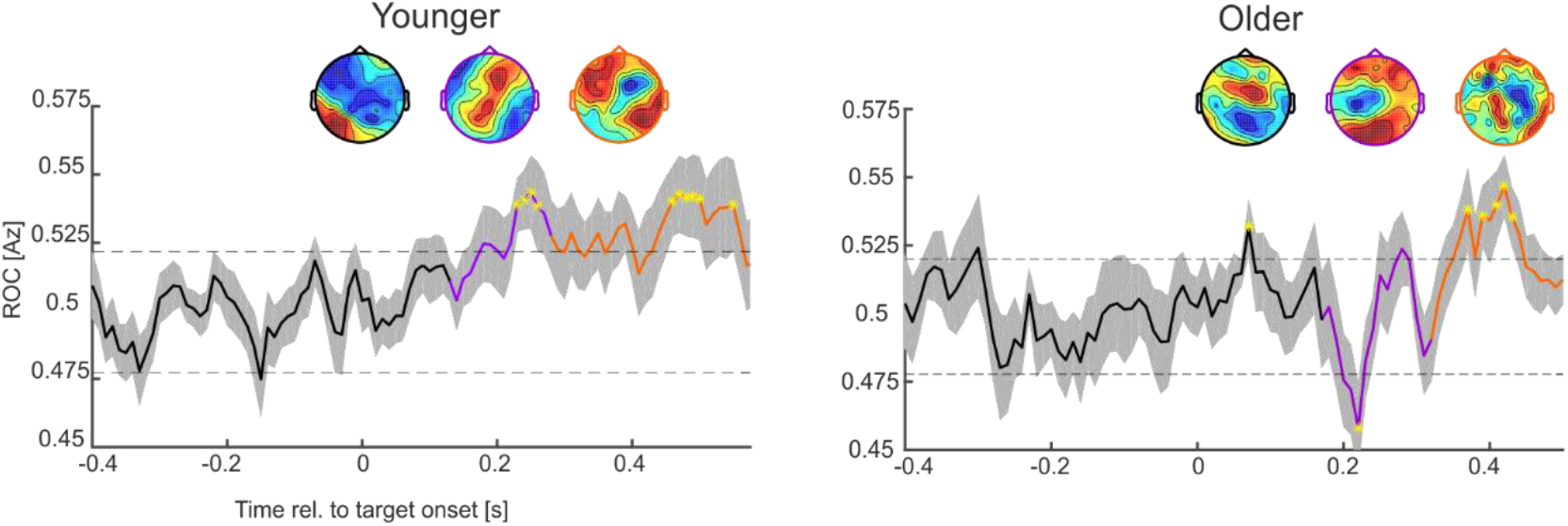
Task-relevant EEG components. A linear classifier based on EEG data in 80ms windows was used to discriminate between the two frequency conditions of interest. The smooth curve reflects group-averaged ROC values (Az) with SEM represented by shaded boundaries. Yellow asterisks highlight projections in which Az reached significance, and the dashed lines represents significance, based on randomisation tests (at p<0.001). Coloured curve segments indicate the k-means clustering of scalp projections derived from the classifier topographies. Clustering revealed three distinct components, each systematically different temporally and topographically. The first cluster (black curve) spanned the epoch in which the stimuli was being presented; the second (purple) cluster comprises early auditory activity; and the third (orange) cluster reflects later decision-making activity. Topographies represent the group-averaged scalp projections of peak Az performance within each cluster.

Using data-driven clustering based on individual subject’s component topographies we extracted three temporally and topographically distinct component-clusters for each group (Kayser et al., 2016). For each of these clusters we derived the respective group-level topographies and classifier performance (Fig. 3 inserts). Importantly, this analysis allowed us to incorporate inter-individual differences in the precise timing of relevant EEG activations, as within each of the three clusters, we selected for each subject the time point at which the respective discriminant component carried maximal information about the stimulus conditions.

The first EEG component spanned a time window encapsulating the majority of the stimulus presentation period (0s to 0.15s) in both younger (0s to 0.12s) and older (0s to 0.16s) participants. Given the overlap with ongoing stimulation, this component was not considered further (see also Kayser et al., 2016). The second component (hereinafter termed the “auditory component”) spanned an early epoch (0.13s to 0.28s in younger, and 0.17s to 0.31s in older adults) and had a central topography in both groups, thus likely reflecting early auditory processing. The third component (hereinafter termed the “decision-making component”) spanned a later trial time epoch (0.28s to 0.5s in younger, and 0.32s to 0.5s in older adults). This late component likely reflects the transition between sensory encoding and conscious perception and was characterized by parieto-frontal contributions (Diaz et al., 2017; Giani et al., 2015; Marti et al., 2015). Both the auditory and decision-making components significantly discriminated between frequency order conditions in both age groups (ROC >0.5; randomization test, p<0.01).

Noteworthy, while the overall topographic sequence of EEG components was the same across groups, the timing of the auditory and decision components was delayed by about 40ms in the older group, which matches the latency shift observed in the ERP P2 component.

### 3.5. Influence of Pre-Target Activity within Auditory Networks

Having derived projections of single-trial task-related activity within meaningful EEG components we computed pre-target oscillatory activity for each component (Fig. 4A). We then used statistical modelling to understand the tri-partite relation between prestimulus activity, the encoding of task-relevant information (as reflected by the EEG component) and behavioural choice (Fig. 4B). Specifically, we statistically tested the relations between power/phase (individually) and choice (model 1); power/phase and sensory evidence (model 2); and sensory evidence and choice (model 3).

**Fig. 4.**
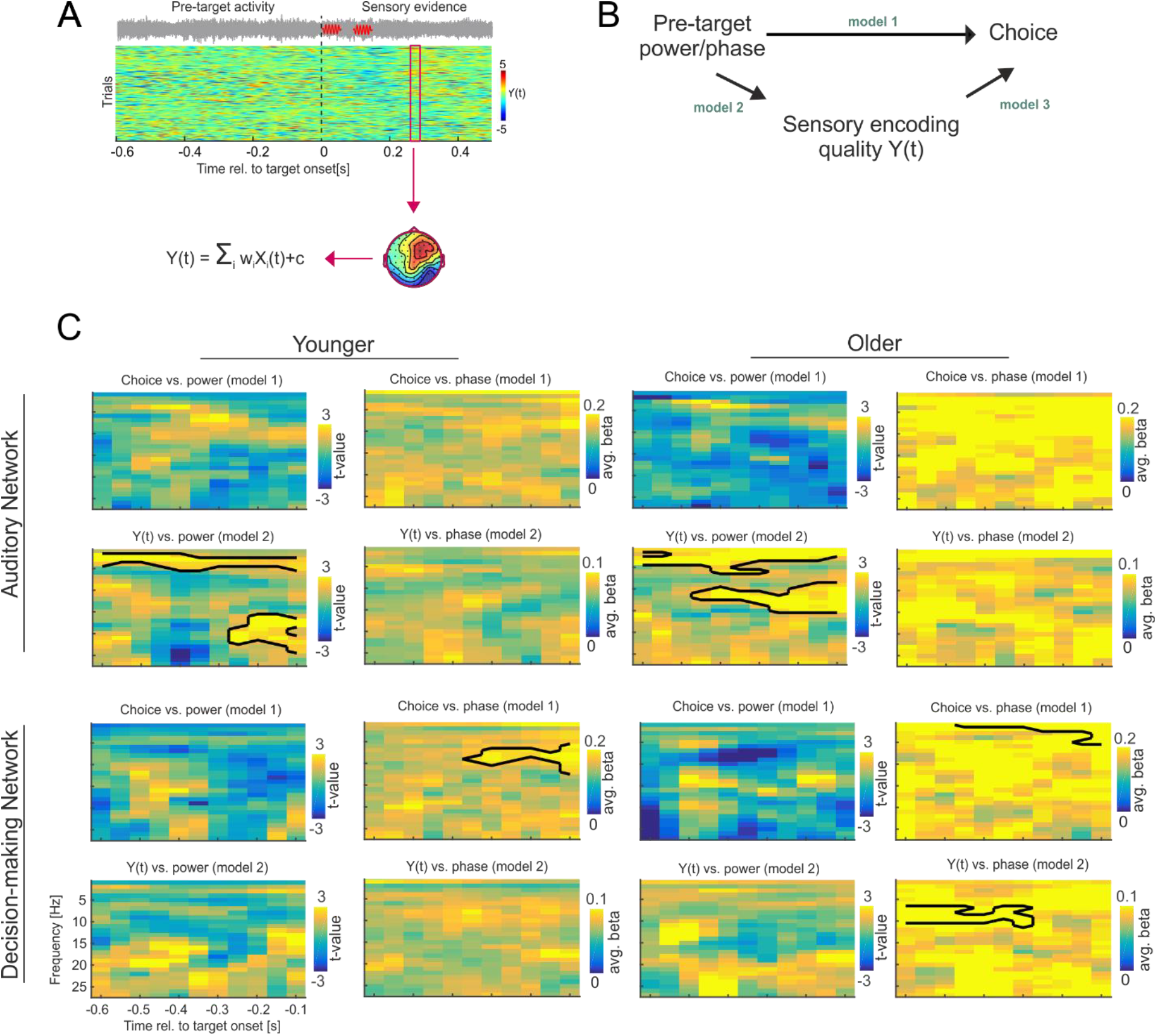
Linear modelling of pre-target activity on sensory evidence and choice within auditory and decision-making networks. (A upper panel) Single trial activity for one participant is shown. The red box highlights a classifier time window. (A lower panel) One-dimensional scalp projections carrying task relevant sensory evidence, *Y*(*t*) are derived from the single trial EEG data, *X*(*t*) and are defined by spatial weights, *w*(*t*), an a constant, c. (B) Models by which pre-stimulus activity could shape perceptual choice. Model 1 tested for influences of power/phase on perceptual choice (i.e. without involving sensory evidence). Model 2 tested for influences of power/phase on Y(t), while model 3 tested for influences of both power/phase and Y(t) on choice. Mediation effects of power/phase on choice via sensory evidence were assessed by comparing models 2 and 3. (C) Group-level regression statistics for models 1 and 2, for both age groups and components. Significant time-frequency clusters are highlighted by black contours (at p<0.05; FDR corrected across models and comparisons).

For the auditory EEG component, we found no significant relation between pre-target power or phase on choice in either group (model 1; at p < 0.05 FDR corrected across models, Fig. 4C). However, there were significant relations between pre-target power and sensory evidence (model 2): in the younger group at low-frequencies (2-6Hz, −0.6s to −0.1s; Tsum = 66, p = 0.001) and the beta band (16-36Hz, −0.3s to −0.1s, Tsum = 77, p = 0.002). The same effects were observed in the older group, albeit at slightly different frequencies: low frequency (2-7Hz, −0.6s to −0.1s, Tsum = 128.6, p < 0.001) and alpha/beta band (10-16Hz, −0.45s to −0.1s, Tsum = 9.6, p =0.001). No relation between phase and sensory evidence was found in either group. The relation between sensory evidence and choice (model 3) was significant in the younger group (t(12) =3.3, p = 0.006) and approached significance in older group (t(11) = 2.1, p = 0.054).

These results could be seen to suggest that pre-stimulus influences emerge systematically at lower frequencies in the older group. However, the existence of a significant cluster at a specific frequency does not demonstrate that this effect is significant only at that specific frequency. We hence used a bootstrap test to directly probe whether the group-level peak frequencies for each cluster differed significantly between groups. This was not the case for either cluster (low frequency cluster: difference in peaks = 0Hz, 95% bootstrap CI [-9,+9] Hz, p=0.199; alpha/beta cluster: difference in peaks = 7Hz, CI [-13,+13] Hz, p=0.173).

### 3.6. Influence of Pre-Target Activity within Decision Networks

Repeating the same comparison for the choice-related EEG component revealed a significant relation between pre-target phase and choice in the younger group around the alpha band (7-14Hz, −0.4s to 0.1s, T_sum_ = 5, p =0.003; Fig. 4C) and at low-frequencies in the older group (1-5Hz, −0.4s to −0.1s, T_sum_ = 4. 5, p = 0.003). Here we found some mild evidence that the respective peak frequencies may differ with age, as the difference was statistically significant (difference in peaks = 7Hz, CI [-8,+9] Hz, p=0.049).

Furthermore, there were no significant relations between power and choice in either group (model 1 for power) and there were no significant relations between power and sensory evidence (model 2). However, in the older group there was a significant relation between alpha phase and evidence (8-12Hz, −0.6s to −0.3s, T_sum_ = 3. 1, p = 0.002), while no such effect was observed in the younger group. Additional mediation analysis revealed no significant mediation effects of phase on choice through evidence in either age group (at p < 0.05), and neither age group showed a significant relation between sensory evidence and choice (younger adults: t(12) = 1.1, p = 0.27; older adults: t(11) = 1.9, p = 0.0741), suggesting that the statistical relation between alpha phase and sensory evidence in the older group reflects a process not directly driving perceptual decisions.

Given that we found some evidence for pre-stimulus influences on choice to emerge at different frequencies in younger and older participants, we also asked whether this difference in peak frequency is related to the observed changes in amplitudes or latencies of the evoked potentials (c.f. Fig. 2). Specifically, we obtained leave-one-out estimates of the group-level peak frequencies for the pre-stimulus effects and EVP amplitudes and latencies in response to background onset. We then used these six EVP characteristics as predictors for the pre-stimulus peak frequencies across the sample of young and old participants. Together the EVP characteristics provided significant predictive power (r^2^ =0.81, F=16.8, p<10^−5^), suggesting that changes in the timing and amplitude of evoked responses are indeed related to the observed changes in relevant pre-stimulus frequencies.

## 4. Discussion

In the current study we investigated the consistency of how pre-stimulus activity influences auditory discrimination performance in young and older participants. In both groups the power of pre-stimulus activity influenced the encoding of sensory evidence in early auditory networks (as read-out by the EEG discriminant component), while the phase influenced choice formation in later-activated fronto-parietal networks.

Importantly, we did not find systematic evidence for a significant difference in the time scales of the involved brain activity between groups for effects arising from auditory cortical networks, while our data revealed a trend for later effects to emerge at slower frequencies in the aging brain. At the same time our data replicate previous findings of a significant age-related slowing of ERP latency, modulations of ERP amplitudes, and a flattening of the spectral profile of EEG activity.

### 4.1. Pre-stimulus influences on perception

Our results confirm previous research showing that pre-stimulus rhythmic activity influences auditory perceptual decisions (Florin, Vuvan, Peretz, & Baillet, 2017;Henry et al., 2014;Henry & Obleser, 2012; Kayser et al., 2016; Ng et al., 2012; Pinheiro et al., 2017; Strauss et al., 2015). These studies, in addition to those investigating such effects in other sensory domains (Hanslmayr et al., 2011; Iemi et al., 2017; Samaha et al., 2017; Samaha and Postle, 2015; VanRullen, 2016), collectively point to a causal role of rhythmic activity in shaping perception regardless of age.

In a previous study focusing on young subjects we dissociated two mechanisms by which pre-stimulus activity influences auditory perception and mapped these onto distinct neural generators. Specifically, we found that low frequency and alpha/beta power shape the encoding of relevant sensory information in early-activated auditory networks, while the phase of alpha band activity directly influences the decision process in later-activated fronto-parietal networks. Here we replicated these results in a group of elderly participants characterized by no or mild hearing loss, in a paradigm where the overall task performance was equated between groups. Thereby the present data lend additional support to the hypothesis that multiple and distinct rhythmic processes control perceptual decisions and suggest that the relevant time scales of neural activity are largely conserved along the life span. This conversation of relevant mechanisms was especially prominent in the shorter-latency auditory-cortical EEG component, despite an overall significant delay of the auditory evoked potential.

### 4.2. Age-related changes in the timing of brain activity

In our data we found systematic age-related differences in the P1-N1-P2 components of auditory evoked responses. Older adults’ P1 and N1 component amplitudes were significantly larger compared to younger adults, yet their P2 peaks were reduced. These findings are consistent with previous reports of age-related changes in AEP amplitude (Anderer et al., 1996; Czigler et al., 1992; Harkrider et al., 2005; Henry et al., 2017; Rufener et al., 2014; Tremblay et al., 2003), which may be attributed to age-related changes at the cellular level (Caspary et al., 2008; de Villers-Sidani et al., 2010; Hughes et al., 2010) or neuronal synchrony (Anderson et al., 2012; Harris and Dubno, 2017). Furthermore, we also found an age-related slowing of the N1 and P2 peak latencies, an effect consistently reported in ageing research (Henry et al., 2017; Tremblay et al., 2004). Such an age-relating slowing of sensory responses may be associated with molecular changes affecting the efficacy of neural signalling (Caspary et al., 2008, 2005, 1995; de Villers-Sidani et al., 2010; Hughes et al., 2010) or changes in white matter/myelination (Lu et al., 2011, 2013; Price et al., 2017; Salat et al., 2005).

We also found that the spectral profile of ongoing EEG activity was significantly flatter in the older participants. This is in line with previous reports, which proposed a mediating role of spectral flattening in cognitive decline (Tran et al., 2016; Voytek et al., 2015), possibly resulting from a decrease in neuronal synchrony (Podvalny et al., 2015; Pozzorini et al., 2013; Voytek and Knight, 2015; Waschke et al., 2017), increases in spontaneous activity (Hong and Rebec, 2012), or changes in the excitation inhibition balance (Caspary et al., 2008; Gao et al., 2017). Our participants passed a cognitive screening assessing a wide variety of cognitive abilities (reasoning, attention, working memory, abstraction, orientation, language), suggesting that the observed changes in spectral slope do not reflect cognitive decline itself but either compensatory mechanisms or basic changes in cellular physiology.

Our main question was whether pre-stimulus influences on perceptual decisions are comparable between young and older participants. While the statistical clusters of significant effects seemed to be systematically shifted towards lower frequencies in the older group, in particular for a direct relation between pre-stimulus phase and choice, direct statistical tests did not provide clear evidence for such an effect. In particular within the early-activated EEG component, which likely reflects neural generators within the auditory cortices, there was no statistical indication of peak frequencies to differ between groups. This could indicate that the processes of early sensory encoding are conceptually conserved with age, despite a slowing of the sensory evoked response.

One possibly is that the present sample size was not sufficient to reveal systematic shifts in the relevant frequencies or that such effects are smaller than the frequency resolution employed here. On the other hand it could also be that the mechanisms and time scales by which pre-stimulus activity shapes sensory encoding remain indeed the same, despite an overall change in the relative amplitude of different frequency bands (Babiloni et al., 2006; Cummins and Finnigan, 2007; Rondina et al., 2016; Vlahou et al., 2014). Support for the latter conclusion comes also from studies demonstrating a similar modulation of alpha band activity by acoustical structure and task demands in young and elderly participants (Erb and Obleser, 2013; Tune et al., 2018; Wostmann et al., 2015), from the comparable alignment of lateralized alpha modulations to acoustic stimuli across age groups (Tune et al., 2018), and from a study demonstrating a similar modulation of behavioural performance by stimulus-entrained delta-band activity in young and older participants (Henry et al., 2017). Furthermore, while many studies confirm age-related changes in the power of individual frequency bands with age, it remains unclear whether the peak frequencies of well-known brain rhythms, such as parietal alpha, change with age (Hong et al., 2015; Klimesch, 1999; McEvoy et al., 2001; Vlahou et al., 2014). In those studies reporting differences the effects are often at the edge of significance (Hong et al., 2015; McEvoy et al., 2001), and a recent study with a larger sample size did not find strong evidence for such changes (Vlahou et al., 2014). The stability of the time scales of prevalent brain rhythms could be one reason for why the pre-stimulus influences on perception remain the same with age, despite an overall slowing of sensory evoked responses.

### 4.3. Alpha activity, cognitive strategies, and aging

Within the later-activated fronto-parietal component the pre-stimulus effects on choice were more variable, and we observed a trend towards a reduction in peak frequencies in the older group. Furthermore, this reduction in peak frequencies was statistically significantly related to changes in the timing and latency of evoked responses between groups. Given that this EEG component likely captures high-level cognitive and decision-making processes our results support the notion that cognitive and decision strategies become more variable with age (McGovern et al., 2017; Sander et al., 2012; Zanto and Gazzaley, 2014). In this context alpha band activity provides an important marker of relevant neural processes. The lateralization of alpha power is indicative of visual-spatial attention (Thut et al., 2012; Wöstmann et al., 2016) and alpha band activity supposedly reflects the selection of sensory information by inhibition (Jensen and Mazaheri, 2010), for example via modulating the excitability of sensory cortices (Iemi et al., 2017; Kayser et al., 2015; Strauss et al., 2015). In auditory perception, the enhancement of alpha activity is often inversely related to signal intelligibility (Becker et al., 2013; McMahon et al., 2016; Obleser et al., 2012; Obleser and Weisz, 2012; Scharinger et al., 2014; Wostmann et al., 2015), possibly reflecting age-related changes in attentional selection (Gazzaley et al., 2005; Henry et al., 2017; Wostmann et al., 2015). Therefore, it may not be surprising that the relation between alpha activity and perceptual decision making differed between groups. The relation between alpha phase and perceptual choice in the later-activated decision component seemed to be systematically shifted towards lower frequencies, albeit we emphasize that the statistical evidence for such a shift was still weak. This frequency difference may reflect distinct decision-making approaches in the two groups, which possibly become slower in the elderly (McGovern et al., 2017). Furthermore, there was also an additional effect of alpha phase on sensory evidence in the decision-related component that was significant only in the older group. While this phase-effect did not directly influence subjects’ choice, and hence did not bear direct influence on behaviour, it could reflect the differential engagement of attentional processes and their impact on the encoding of sensory information in younger and older listeners (Henry et al., 2017; Wostmann et al., 2015).

## 5. Conclusion

The present data demonstrate conceptually similar influences of rhythmic pre-stimulus activity on sensory encoding in young and older healthy listeners. This consistency in pre-stimulus effects arises largely despite systematic changes in the overall spectral profile of EEG activity and a general slowing of auditory evoked responses in the older participants, raising questions as to how these two processes are biophysically related. At the same time, we observed a trend towards a distinct influence of the timing of alpha and delta/theta band activity in later-activated EEG components with age, which calls for a more systematic assessment of the relation between rhythmic brain activity, sensory encoding and cognitive strategies in aging.

## Acknowledgments

This work was supported by the UK Biotechnology and Biological Sciences Research Council (BBSRC; BB/L027534/1). S.W.M. is supported by a studentship from the UK Economic and Social Research Council (ESRC; 1820770). C.K. is supported by the European Research Council (ERC-2014-CoG; Grant 646657).

